# 3D UNet with GAN discriminator for robust localisation of the fetal brain and trunk in MRI with partial coverage of the fetal body

**DOI:** 10.1101/2021.06.23.449574

**Authors:** Alena Uus, Irina Grigorescu, Milou van Poppel, Emer Hughes, Johannes Steinweg, Thomas Roberts, David Lloyd, Kuberan Pushparajah, Maria Deprez

## Abstract

In fetal MRI, automated localisation of the fetal brain or trunk is a prerequisite for motion correction methods. However, the existing CNN-based solutions are prone to errors and may require manual editing. In this work, we propose to combine a multi-label 3D UNet with a GAN discriminator for localisation of both fetal brain and trunk in fetal MRI stacks. The proposed method is robust for datasets with both full and partial coverage of the fetal body.

## 1 Introduction

In clinical practice, fetal MRI acquisition protocols commonly include stacks of single shot fast spin echo (ssFSE) *T*_2_-weighted slices acquired under different multiple orientations, that may cover the whole uterus, or only the fetal brain or trunk separately [1]. Fetal motion is generally addressed by slice-to-volume registration (SVR) methods that have been widely employed for 3D reconstruction of the fetal brain [2, 3, 4, 5] and thorax [6]. The recently proposed deformable SVR (DSVR) allows 3D reconstruction of the whole body [7]. However, SVR-based methods require localisation of the regions of interest (ROI) as an input. Recently, a series of CNN segmentation-based solutions were proposed for fetal brain localisation and automation of SVR reconstruction. These works employed 2D UNet [8], 2D P-net [4] or a 3D V-net [9]. The output 3D masks were then refined using morphological operations. In [10], a 3D UNet was used for localisation of the whole fetus in EPI stacks. However, 2D segmentations often lead to errors due to insufficient context information and 3D networks fail when the object is not present in a stack due to partial coverage. Therefore, the existing localisation pipelines may require manual editing.

### 1.1 Contributions

In this work, we propose a technique for automated robust simultaneous localisation of both fetal brain and trunk in stacks with either full or partial coverage of the uterus. It is based on integration of a 3D UNet with multiple labels and a GAN discriminator module that minimises the erroneously segmented regions.

## 2 Methodology

### 2.1 Datasets, acquisition and preprocessing

The data used in this work includes T2 ssFSE MR datasets of 43 fetal subjects at 28-33 weeks GA acquired at Evelina Children’s Hospital, London as a part of fetal CMR practice. Each of the datasets includes 10-13 stacks acquired under different orientations and with different coverage of the fetal body, covering either whole uterus, or only fetal brain or trunk ROIs.

For each of the datasets, the 3D masks covering the approximate brain, body and uterus ROIs were manually drawn on one of the stacks transferred to the rest using label propagation. The stacks and the corresponding masks were resampled with padding to 128×128×128 voxel grid size with the resulting 3-5 mm varying isotropic resolution.

### 2.2 Proposed pipeline for fetal localisation

Our proposed fetal localisation pipeline uses the paradigm of adversarial training, by including a discriminator network to fine tune the predicted labels obtained from the segmentation module. For the latter, we employ a 3D UNet [11] architecture with 5 encoding-decoding branches with 32, 64, 128, 256 and 512 channels, respectively. Each encoder block consists of 2 repeated blocks of 3 × 3 × 3 convolutions (with a stride of 1), instance normalisation [12] and LeakyReLU activations (see 3D UNet layer in Figure 1). The first two down-sampling blocks contains a 2 × 2 × 2 average pooling layers, while the others use 2 × 2 × 2 max pooling layers. The decoder blocks have a similar architecture as the encoder blocks, followed by upsampling layers. The model outputs an N-channel 3D image, corresponding to our N classes: background, uterus, fetal brain and fetal trunk.

**Figure 1:**
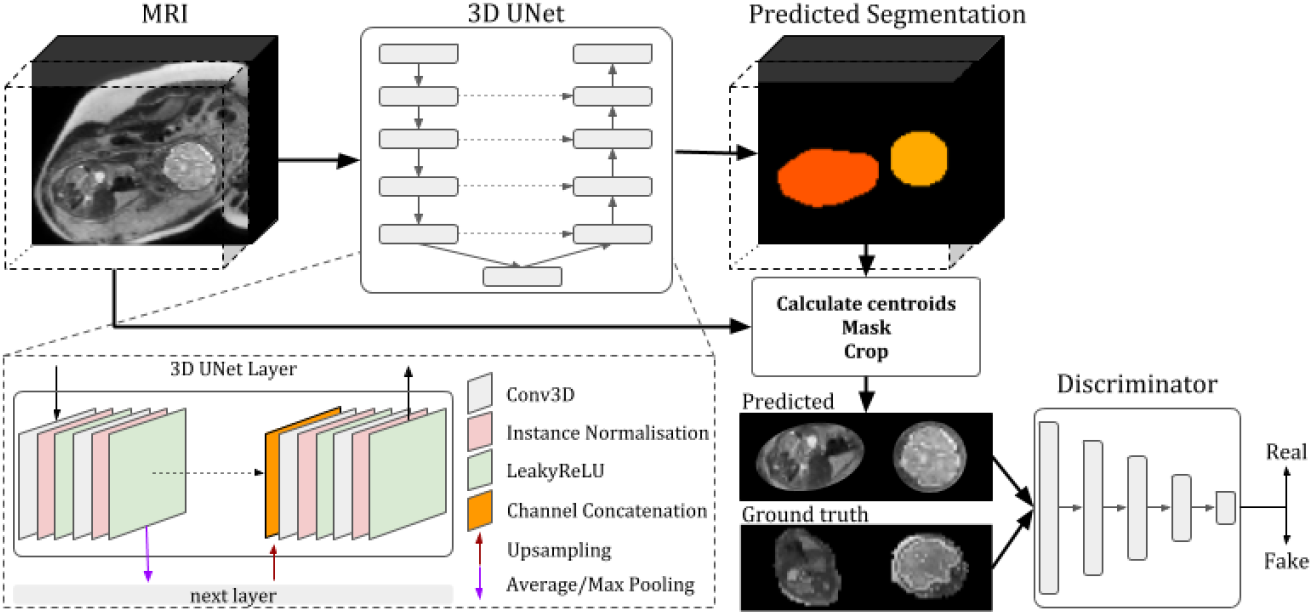
The proposed pipeline for localisation of fetal body and brain in 3D MRI stacks.

The discriminator network used in our pipeline (see Figure 1) is a PatchGAN discriminator as proposed in [13]. It is made up of repeated blocks of 3D convolutions (with 64, 128, 256 and 512 channels), instance normalisation layers [12] and LeakyReLU activations.

The segmentation network is trained by minimizing a generalised Dice loss (see Equation 1) [14] using the Adam optimizer with the default parameters (*β*_1_ = 0.9 and *β*_2_ = 0.999).

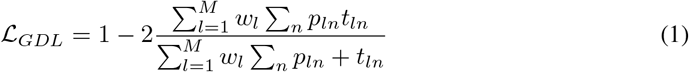

where *w_l_* = 1/(Σ_*n*_ *t*_*ln*_)^2^ is the weight of the l^*th*^ tissue type, *p*_*ln*_ is the predicted probabilistic map of the l^*th*^ tissue type at voxel n, *t*_*ln*_ is the target label map of the l^*th*^ tissue type at voxel n, and M is the number of tissue classes. The discriminator network is trained using the Adam optimiser with *β*_1_ = 0.5 and *β*_2_ = 0.999. Both networks are trained with a learning rate of 2 · 10^−3^ with a linearly decaying scheduler. When training the discriminator, we use the Least Squares GAN [15] loss function: 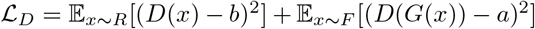 where *a* represents the label for fake images and *b* is the label for real images. The generator and the segmentation network were trained together using 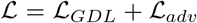, where 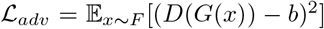. The networks were implemented in PyTorch with TorchIO library [16] for data augmentation. The code is available online as a part of SVRTK toolbox^2^.

## 3 Experiments and Results

In order to assess the performance of the proposed 3D UNet+GAN localisation approach, we compare it to a classical 3D UNet with multiple (uterus+brain+trunk) and single (brain or trunk only) labels. We used 32 datasets (381 3D MRI stacks) for training, 3 for validation (36 3D MRI stacks) and 8 for testing (100 3D MRI stacks). All methods were trained for 300 epochs with the same input datasets and augmentations settings (affine transformations, motion and spike artefacts). In addition, for all methods, we post-processed the segmentations by excluding extremely small components.

Typical examples of localisation outputs in stacks with full and partial field of view (FoV) are demonstrated in Figure 2. In general, 3D UNet based localisation showed high performance under the various levels of motion corruption of stacks (Figure 2.A) but sometimes it led to errors if the structure of interest was not present in the FoV (Figure 2.B). This was resolved by the GAN discriminator.

**Figure 2:**
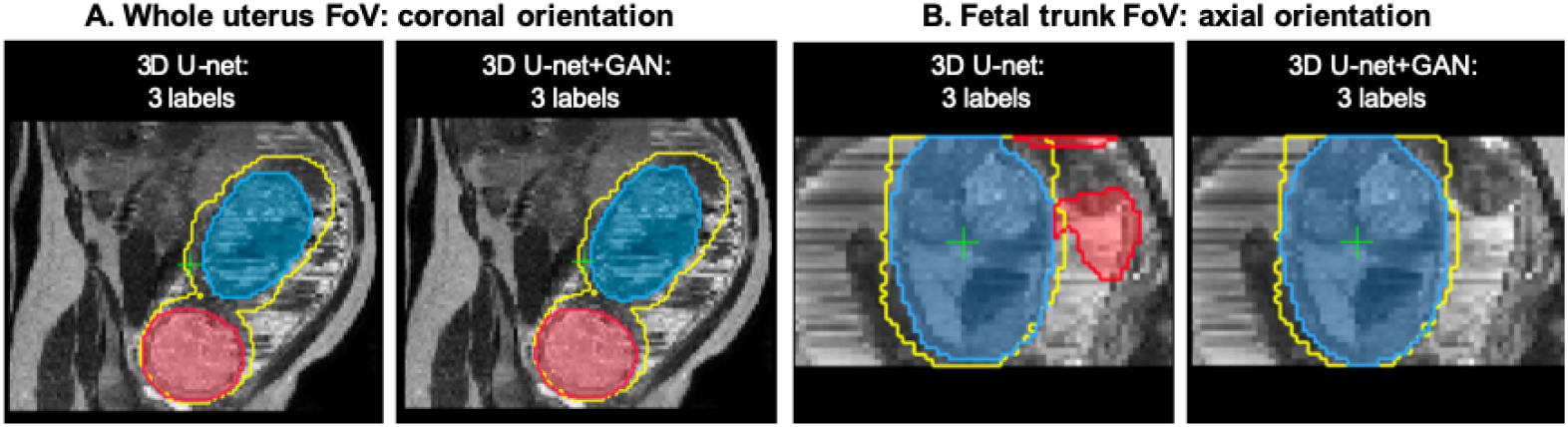
An example of fetal brain (red) and trunk (blue) localisation in stacks with the whole uterus (A) and partial fetal body (B) FoVs. The GT labels are shown as yellow contours.

We evaluated localisation performance on 8 test datasets with 100 stacks in total, based on the centroid distance (*d*[*mm*]), the false positive rate (*FPR*) and the manually graded localisation quality scores (*LQS*) defined as *{0-incorrect; 1-partially correct; 2-correct}*, see Table 1. The high centroid distances and FPR values in the baseline 3D UNet results correspond to wrong localisation outputs, while differences in small values are not informative, because the ground truth (GT) masks are not precise. The UNet+GAN correctly localised brain and trunk in all datasets, while for 3D UNet the brain localisation failed in approximately 10% of the cases. Trunk localisation using 3D UNet with a single label failed in 16% of the cases. There were no failures for fetal trunk for 3D UNet with 3 labels, but the mean *LQS* demonstrated that incorporating GAN discriminator resulted in qualitatively better localisation of both trunk and brain.

**Table 1:**
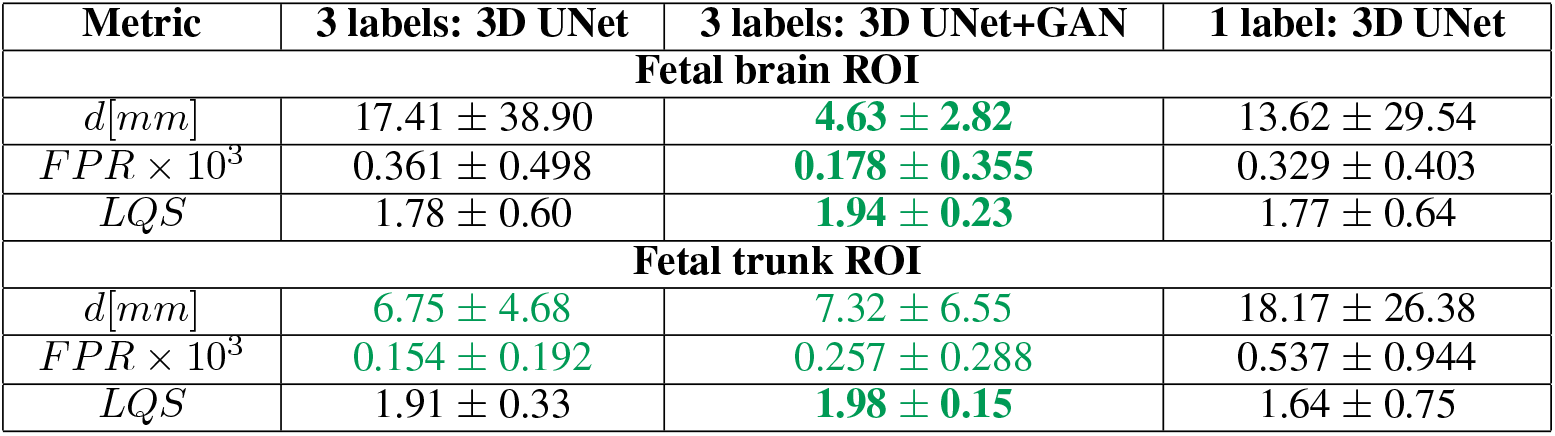
Evaluation of the fetal brain and trunk localisation quality of the proposed 3D UNet+GAN in comparison to the 3D UNet with multiple or single labels. The *FPR* and *d* results are statistically significant with *p* < 0.01 apart from the difference between the 3 label networks for the trunk ROI.

## 4 Conclusions

This work presented a solution for robust automated 3D localisation of fetal brain and trunk in MRI stacks with partial coverage of fetal body. It is based on a 3D UNet with multiple labels extended with a GAN discriminator module. The results demonstrated that 3D UNet+GAN outperforms the 3D UNet segmentation-only approach by preventing the errors when the brain or trunk are not within the FoV. In future, we will focus on optimisation for the wider GA range, various acquisition parameters and integration into the SVR and DSVR reconstruction pipelines.

### Broader Impact

Motion correction methods for fetal MRI already showed a high potential to reduce scan times and improve the information content available for diagnosis. And, in general, the existing various SVR-based pipelines for fetal MRI provide operational solutions acceptable for research purposes. However, automation as well as the fast reconstruction with controlled quality are the key factors required for deployment of SVR methods in the clinical settings.

This work will contribute to development of a robust reconstruction pipeline for automated and fast motion correction of brain, body and placenta that will not require manual input and will provide a stable quality solution. The main limitation of the current study is the small number of subjects and a specific acquisition protocol that might not fully reflect a broad range of different aspects of fetal MRI acquisition practices used at different hospitals. This will be addressed in future work.

## Acknowledgments

We thank everyone who was involved in acquisition of the datasets and all participating mothers.

This work was supported by the Rosetrees Trust (A2725), the Wellcome/EPSRC Centre for Medical Engineering at King’s College London (WT 203148/Z/16/Z), the NIHR Clinical Research Facility (CRF) at Guy’s and St Thomas’ and by the National Institute for Health Research Biomedical Research Centre based at Guy’s and St Thomas’ NHS Foundation Trust and King’s College London. The views expressed are those of the authors and not necessarily those of the NHS, the NIHR or the Department of Health.

## Footnotes

2 SVRTK fetal segmentation: https://github.com/SVRTK/Segmentation_FetalMRI

